# Naturally occurring fluorescence protects the eutardigrade *Paramacrobiotus* sp. from ultraviolet radiation

**DOI:** 10.1101/851535

**Authors:** Harikumar R Suma, Swathi Prakash, Debasish Giri, Govindasamy Mugesh, Sandeep M. Eswarappa

**Author notes:** Correspondence: Sandeep M Eswarappa, Assistant Professor, Department of Biochemistry, Indian Institute of Science, Bengaluru 560012, Karnataka, India.

## Abstract

Naturally occurring fluorescence has been observed in multiple species ranging from bacteria to birds. In macroscopic animals such as birds and fishes, fluorescence provides a visual communication signal. However, the functional significance of this phenomenon is not known in most cases. Though photoprotection is attributed to fluorescence under ultraviolet (UV) light in some organisms, it lacks direct experimental evidence. Here, we have identified a new species of eutardigrade belonging to the genus *Paramacrobiotus*, which exhibits fluorescence under UV light. Using a natural variant of the same species that lacks fluorescence, we show that the fluorescence confers tolerance to lethal UV radiation. Remarkably, we could transfer this property to UV-sensitive *Hypsibius exemplaris*, another eutardigrade, and also to *C. elegans*, a nematode. Using high performance liquid chromatography (HPLC) we isolated the fluorescent compound from *Paramacrobiotus* sp. This compound has excitation maxima (λ_ex_) at 370 nm and emission maxima (λ_em_) at 420-430 nm. We propose that *Paramacrobiotus* sp. uses a fluorescent shield that absorbs harmful UV radiation, and emits harmless blue light, thereby protecting itself from the lethal effects of UV radiation.

**Summary statement:** Tardigrades are well known for their tolerance to extreme environmental conditions. In this study, we have identified a new tardigrade species that employs a fluorescent shield to protect itself from the germicidal ultra violet radiation.

## Introduction

Tardigrades (water bears) are microscopic (0.5 to 1 mm) limno-terrestrial animals with four pairs of legs. More than a thousand species have been reported worldwide from various habitats (Guidetti and Bertolani, 2005). *Hypsibius exemplaris*, *Ramazzottius varieornatus, Richtersius coronifer*, *Milnesium tardigradum* and *Paramacrobiotus richtersi* are some of the well-studied tardigrade species. Phylum Tardigrada comes under the superphylum Ecdyzoa along with its sister phyla: Arthropoda, Nematoda and Onychophora (Borner et al., 2014). An analysis based on the *Hox* genes revealed that the entire tardigrade body is homologous to just the head segment of arthropods (Smith et al., 2016).

Tardigrades are known for their tolerance to extreme environmental conditions. Among them, their ability to tolerate complete desiccation is well known (Goldstein and Blaxter, 2002). This is achieved by a reversible process called anhydrobiosis where tardigrades shrink their size and slow-down their metabolism to reach a physiological state called ‘tun’. Tardigrades are also known to tolerate other physical stresses including extreme temperature, pressure, ionizing radiations, oxidative stress and osmotic variations (Altiero et al., 2011; Guidetti et al., 2011; Hashimoto et al., 2016; Hengherr et al., 2009; Horikawa et al., 2013; Jonsson et al., 2005; Ono et al., 2008; Rizzo et al., 2010; Tsujimoto et al., 2016). These harsh conditions are lethal to most of the animals. Certain tardigrade species have even survived the exposure to space vacuum and galactic radiations in low Earth orbit conditions (Jonsson et al., 2008).

Unfortunately, the molecular and cellular mechanisms behind the extraordinary stress tolerance of tardigrades are poorly understood. Of late, there is an increase in the molecular studies that focus on the stress tolerance of tardigrades (Harikumar and Eswarappa, 2017). A recent analysis of the genome of the tardigrade *Ramazzottius varieornatus* revealed several potential mechanisms behind its extreme radiotolerance. It lacks the genes that promote stress-induced damage and there is a selective expansion of gene families responsible for decreasing various stress-induced damages. This tardigrade has also evolved a unique protein called damage suppressor protein (Dsup), which can impart partial radiotolerance to cultured mammalian cells (Hashimoto et al., 2016). Another study showed that tardigrade-specific intrinsically disordered proteins (TDPs) are required for desiccation tolerance in *Hypsibius exemplaris* (Boothby et al., 2017). Expression of these proteins is increased during desiccation and they form non-crystalline amorphous solids. This vitrification process is implicated in the desiccation tolerance of tardigrades.

Though fluorescence is abundant in marine organisms, it is not very common in terrestrial animals. Fluorescence has been reported in parrots, scorpions, chameleons, frogs, and nematodes (Arnold et al., 2002; Coburn et al., 2013; Lagorio et al., 2015; Protzel et al., 2018; Taboada et al., 2017). Functional significance of this phenomenon is unclear although visual signalling towards potential mates has been attributed in case of parrots (Arnold et al., 2002). Fluorescence has been reported in tardigrades also, but its function is unknown (Perry et al., 2015).

Here we show that an eutardigrade *Paramacrobiotus* sp. exhibits tolerance to germicidal ultraviolet (UV) radiation up to one hour. These tardigrades also show fluorescence under UV light. We demonstrate that this phenomenon contributes to their exceptional tolerance to UV radiation.

## Materials and methods

### Tardigrade culture

*Paramacrobiotus* sp. was isolated from moss samples grown on a concrete wall inside Indian Institute of Science campus (13° 0′ 55″ N and 77° 33′ 57″ E), Bangalore, India. The concrete wall was well lit by direct sunlight most of the days in a year. Moss samples were kept in a 90 mm petri dish immersed in reverse osmosis (RO) water for 12 h. Samples were gently tapped to facilitate the dislodgement of tardigrades to the water in the petri dish, which were then visualized under an upright light microscope (CXL Plus, LABOMED, USA). Once spotted, tardigrades were taken out using a pipette, rinsed in RO water and placed in an embryo culture dish. They were cultured in KCM solution (7 mg KCl, 8 mg CaCl_2_, and 8 mg MgSO_4_.7H_2_O in 1 L of water) in 35 mm petri dishes coated with 2% low EEO agarose (Lonza, Switzerland) at 20° C (Suzuki, 2003). Cultures were kept in dark and *Caenorhabditis elegans* and rotifers were provided as food source. *Hypsibius exemplaris* (previously known as *Hypsibius dujardini Z151*) was a kind gift from Prof. Bob Goldstein University of North Carolina at Chapel Hill, USA. They were cultured as described previously in Chalkley’s medium with algae (*Chlorococcum* sp.) as food source (Gabriel et al., 2007). Prior to experiments, tardigrades were washed thoroughly using RO water and starved for 24 h in RO water containing Ampicillin (0.5 mg/ml). *C. elegans* (Bristol N2 strain) was a kind gift from Dr. Varsha Singh, Indian Institute of Science, Bengaluru. Worms were maintained in Nematode Growth Medium (NGM) with *E. coli* OP50, as the food source.

### Identification of tardigrades by sequencing

Genomic DNA was extracted from a single individual of the newly isolated tardigrade by using a rapid salt and ethanol precipitation as described before (Cesari et al., 2009). A single tardigrade was homogenised in 100 µl of TNES extraction buffer (50 mM Tris pH 7.8, 400 mM NaCl, 20 mM EDTA, 0.5% SDS) followed by proteinase K (0.01 µg/µl) treatment for 2 h at 55° C. To precipitate proteins, 1M NaCl was added and the mixture was centrifuged at 18,000 g for 5 min and the supernatant was collected. Equal volumes of phenol-chloroform-isoamyl alcohol was added to the supernatant and centrifuged at 20,000 g for 5 min. Genomic DNA was precipitated using 70% ethanol and 0.3 M sodium acetate and resuspended in 20 µl of nuclease-free water. PCR reactions were carried out using 200 ng of genomic DNA and universal primers for *COI* (Mitochondrial cytochrome oxidase 1) gene and ITS2 (nuclear Internal Transcribed spacer 2) region as described previously (Folmer et al., 1994; White et al., 1990). The PCR product was subjected to Sanger sequencing. Obtained sequence was subjected to BLAST analysis to identify the closest tardigrade species.

Primers for *COI*: 5′-GGTCAACAAATCATAAAGATATTGG-3′

5′-TAAACTTCAGGGTGACCAAAAAATCA-3′

Primers for *ITS2*: 5′-GCATCGATGAAGAACGCAGC-3′

5′-TCCTCCGCTTATTGATATGC-3′

### Phylogenetic analysis

The COI gene (encodes Cytochrome C oxidase subunit I) sequences of various tardigrade species belonging to the following families were retrieved from NCBI (species name and accession number are given in parenthesis): Macrobiotidae (*Paramacrobiotus richtersi* group 2 from Kenya [EU244598], *Paramacrobiotus richtersi* group 3 from Kenya [EU244599], *Paramacrobiotus richtersi* isolate CLARE_ISLAND-3 [MK040994], *Paramacrobiotus richtersi* isolate CLARE_ISLAND-2 [MK040993], *Macrobiotus pallarii* isolate Tar407 [FJ435807], *Paramacrobiotus lachowskae* [MF568534], *Macrobiotus papei* [MH057763], *Macrobiotus macrocalix* isolate PL.110 [MH057767], *Macrobiotus polypiformis* haplotype 1 [KX810011], *Macrobiotus hannae* strain PL.010 [MH057764], *Macrobiotus shonaicus* haplotype 1 [MG757136]), Hypsibiidae (*Hypsibius exemplaris* strain Sciento Z151 [MG818724], *Ramazzottius varieornatus* [MG432813], *Hypsibius klebelsbergi* voucher HD070 [KT901832],), Milnesiidae (*Milnesium tardigradum* from Japan [EU244604], *Milnesium variefidum* [KT951663], *Milnesium berladnicorum* [KT951659]), and Echiniscidae (*Echiniscus testudo* from Egypt [EU244601], *Echiniscus cf. testudo* Et-Ladakh-1 [EF620367], *Echiniscus blumi* haplotype 1 [EU046090]). The COI sequence of newly identified tardigrade species was obtained by sanger sequencing as described above.

Phylogenetic tree was constructed using MEGA X software (ver. 10.1.5). The COI sequences were aligned using ClustalW option in MEGA X software. All the sequences were trimmed to make them of uniform length. Maximum Likelihood analysis was performed with 1000 bootstrap replicates using a General Time Reversible (GTR) + G+I model.

### Microscopy

Tardigrades and their eggs were placed on a glass slide with two drops of saline (0.9%) solution. Wet mounts were prepared by gently placing a coverslip over the saline drop without damaging the tardigrades. Excess fluid was removed using a lint-free tissue paper. Bright field images were captured using Axio Scope upright light microscope (Carl Zeiss, Germany). DIC (Differential interference contrast) and fluorescence images were obtained using Axio Observer.Z1 inverted fluorescence microscope (Carl Zeiss, Germany) equipped with an HBO 100 lamp. Band pass filters were used for excitation and emission (G365 and BP 445/50 (Carl Zeiss)). DAPI channel was used to capture fluorescence images with exposure time constant at 1 s during the image acquisition. The acquired images were analyzed using Axio Vision imaging software (version: 4.8.2.0; Carl Zeiss, Germany).

### UV Irradiation

Three groups of ten individuals from both tardigrade species were taken in 35 mm petri dishes coated with 2% low EEO agarose and excess water was removed. The animals remained hydrated for the duration of experiment as they were in contact with the moist agarose surface (Horikawa et al., 2013). They were immediately exposed to UV radiation (peak wavelength 253 nm) emanating from a germicidal lamp (LT-T8 30W/UV-C HRA, Narva, Germany). The irradiance of the beam was 0.111 mW/cm^2^ as measured by an UV radiation meter (UVITEC, UK). The UV dose was calculated according to the formula 1 (mW/cm^2^)sec = 1 mJ/cm^2^ as described previously (Horikawa et al., 2013). Tardigrades were exposed to multiple doses (0.6 kJ/m^2^, 1 kJ/m^2^, 2 kJ/m^2^, 4 kJ/m^2^ and 8 kJ/m^2^) of UV light by varying the duration of exposure. After the exposure, samples were transferred to fresh 2% agarose-coated 35 mm petri dishes and cultured as described above. They were monitored daily under a light microscope for a period of 30 days. Any eggs laid were collected and cultured separately to check their hatchability.

To test if the UV resistant property of *Paramacrobiotus* sp. can be transferred to UV sensitive *H. exemplaris* or *C. elegans*, three hundred individuals of *Paramacrobiotus* sp. were homogenized in 120 μl of water using a mechanical tissue grinder. This was followed by centrifugation of the lysate at 20,000 g for 15 min. The supernatant showed fluorescence under UV light (wavelength 254 nm and 365 nm) and 40 μl of it was added in a well of a 96-well plate. 40 μl of sterile water was used as control. Twenty individuals of *H. exemplaris* or 50 individuals of *C. elegans* were added in those wells, and exposed to UV-C radiation (1 kJ/m^2^) as described above. They were monitored every day after the treatment.

### Methanol extraction of tardigrades

Five hundred individuals of *Paramacrobiotus* sp. were transferred into a 1.5 ml tube containing 200 µl methanol. The tube was subjected to a freeze-thaw cycle in liquid nitrogen followed by mechanical homogenization using a tissue grinder. The lysate was centrifuged at 20,000 g for 5 min and the supernatant was collected. This process was repeated twice and the supernatants were pooled. As a control, 1000 individuals of *H. exemplaris* were subjected to the same extraction procedure and the supernatant was collected. The extracts were then visualized under a UV lamp (GeNei^TM^, India) to observe the fluorescence (excitation wavelengths were: 254 nm and 365 nm).

### High Performance Liquid Chromatography (HPLC)

100 μl of methanolic extract of tardigrades was injected into Waters HPLC system with a reverse phase column (Hibar^®^ RT C-18 column; 4.6×250 mm, particle size: 5 µm) using an auto sampler. A gradient of acetonitrile/water was used as the mobile phase at a flow rate of 1 ml/min. A PDA detector was used to detect absorbance at 350 nm. The fraction corresponding to the unique peak around 5.5 to 6 min in *Paramacrobiotus* sp. extract was collected from the flow-through. After confirming the fluorescence of this fraction, it was lyophilized and re-suspended in methanol. Fluorescence properties were investigated using a spectrofluorometer (FP-6300, Jasco, USA). As a control the HPLC fraction of *H. exemplaris* was used. Excitation profile was obtained by varying the emission wavelength between 400 nm and 460 nm. Emission profile was obtained by varying the excitation wavelength between 300 nm and 370 nm. The spectral profiles were corrected by subtracting the values of methanol.

## Results

### The new tardigrade isolate belongs to the genus *Paramacrobiotus*

We identified a new species of tardigrade from moss samples growing on a concrete wall at Indian Institute of Science, India (13° 0′ 55″ N and 77° 33′ 57″ E; see Methods for details). To identify the species, we sequenced the *COI* (Mitochondrial cytochrome oxidase 1) gene and *ITS-2* (nuclear Internal Transcribed spacer 2) region from a single individual using universal primers as described previously (Folmer et al., 1994; White et al., 1990). BLAST analysis of the obtained sequences against all available nucleotide sequences in NCBI revealed their high similarity to the sequences of *COI* (81.62% identity) and *ITS-2* (85.96% identity) from the eutardigrade *Paramacrobiotus richtersi*. Phylogenetic analysis based on these sequences revealed that the newly identified tardigrades belong to the genus *Paramacrobiotus* under the family Macrobiotidae, class Eutardigarda and phylum Tardigrada (Fig 1A and Table S1, S2). Henceforth the new tardigrade species will be referred to as *Paramacrobiotus* sp.

**Figure 1.**
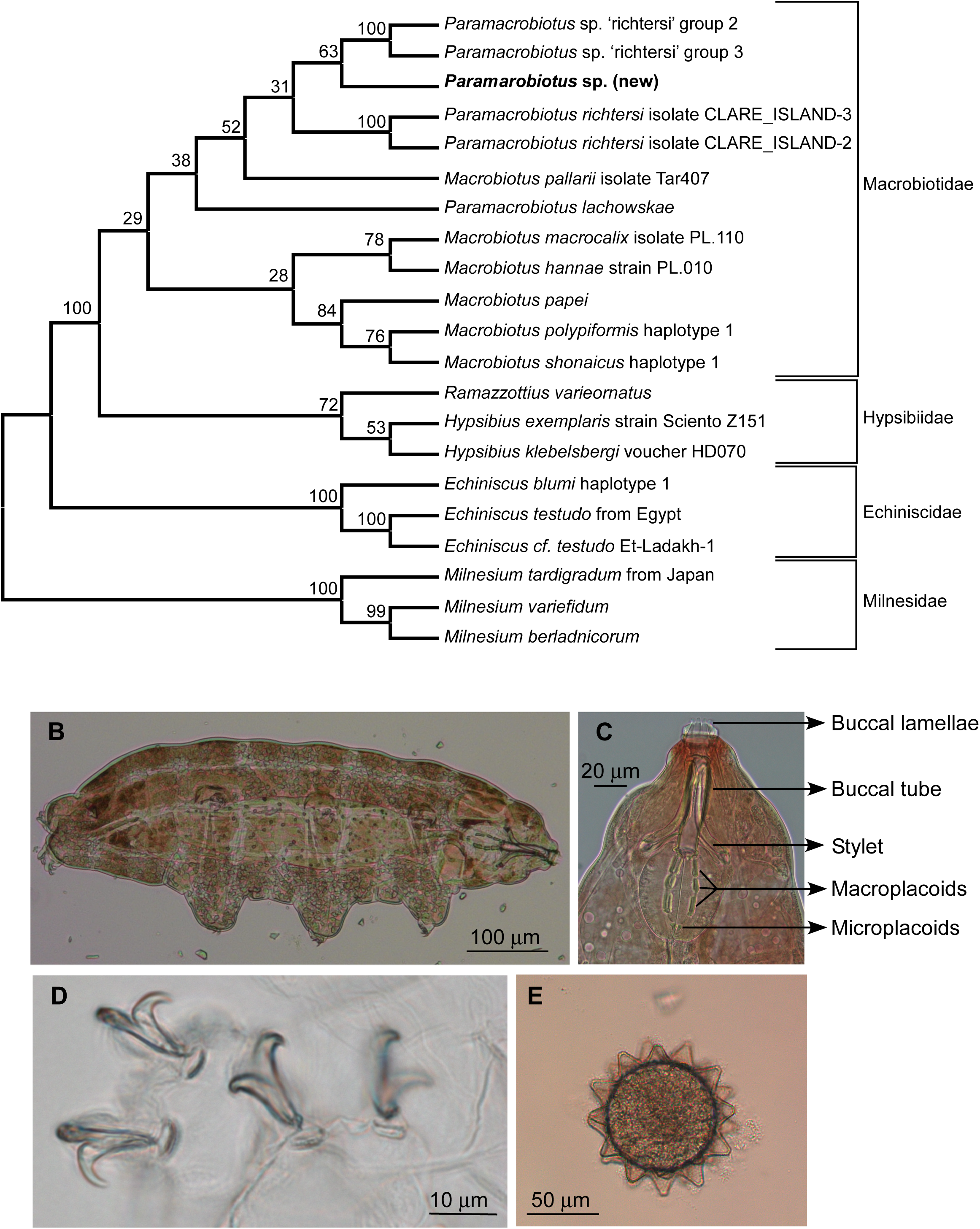
Newly isolated tardigrade species belongs to the genus *Paramacrobiotus*. (A) Maximum likelihood phylogenetic tree based on *COI* sequence of various tardigrades. The tree was constructed using MEGA X software. Newly identified species is shown in bold. *COI* sequences of other tardigrade species were taken from NCBI. Numbers show bootstrap values. (B-E) Morphological features of the new species. The entire body with reddish-brown pigmentation (B), buccopharyngeal (feeding) apparatus (C), hind legs with two Y-shaped double claws (D), and ornamented eggs with surface projections (E) are shown.

These tardigrades were about 600 µm long and had reddish-brown pigmentation (Fig 1B). Close examination of the buccopharyngeal (feeding) apparatus of the new isolate showed a cylindrical buccal tube (mouth tube) attached to an oval shaped pharynx. The mouth opening was surrounded by buccal lamellae in a circular arrangement mounted on the buccal crown. The feeding apparatus had anterior macroplacoids and posterior microplacoids with piercing stylets in a bent position (Fig 1C). Hind legs had two Y-shaped double claws (Fig 1D). The eggs laid by this tardigrade were spherical and ornamented with conical surface projections. They were deposited outside the moulted cuticle (Fig 1E). All these morphological features are characteristic of the family Macrobiotidae to which the genus *Paramacrobiotus* belongs (Guidetti et al., 2012; Schill, 2018).

### *Paramacrobiotus* sp. shows tolerance to UV radiation

Since tardigrades are known for their tolerance to extreme conditions, we exposed the *Paramacrobiotus* sp. to multiple physical stresses. We observed that these tardigrades were particularly resistant to UV radiation. All of the *Paramacrobiotus* sp. survived 10 min exposure (corresponds to 0.66 kJ/m^2^) to germicidal UV radiation, whereas all of the *Hypsibius exemplaris*, another eutardigarde species, died within minutes after the same treatment (Fig 2A). Furthermore, 60% of *Paramacrobiotus* sp. survived 1 h exposure to UV radiation (corresponds to 4 kJ/m^2^) beyond 30 days (Fig 2B, C and D). The survived tardigrades were observed daily; they laid eggs that hatched to normal individuals. This was observed for two generations showing that UV exposure did not affect their survival or their reproductive ability.

**Figure 2.**
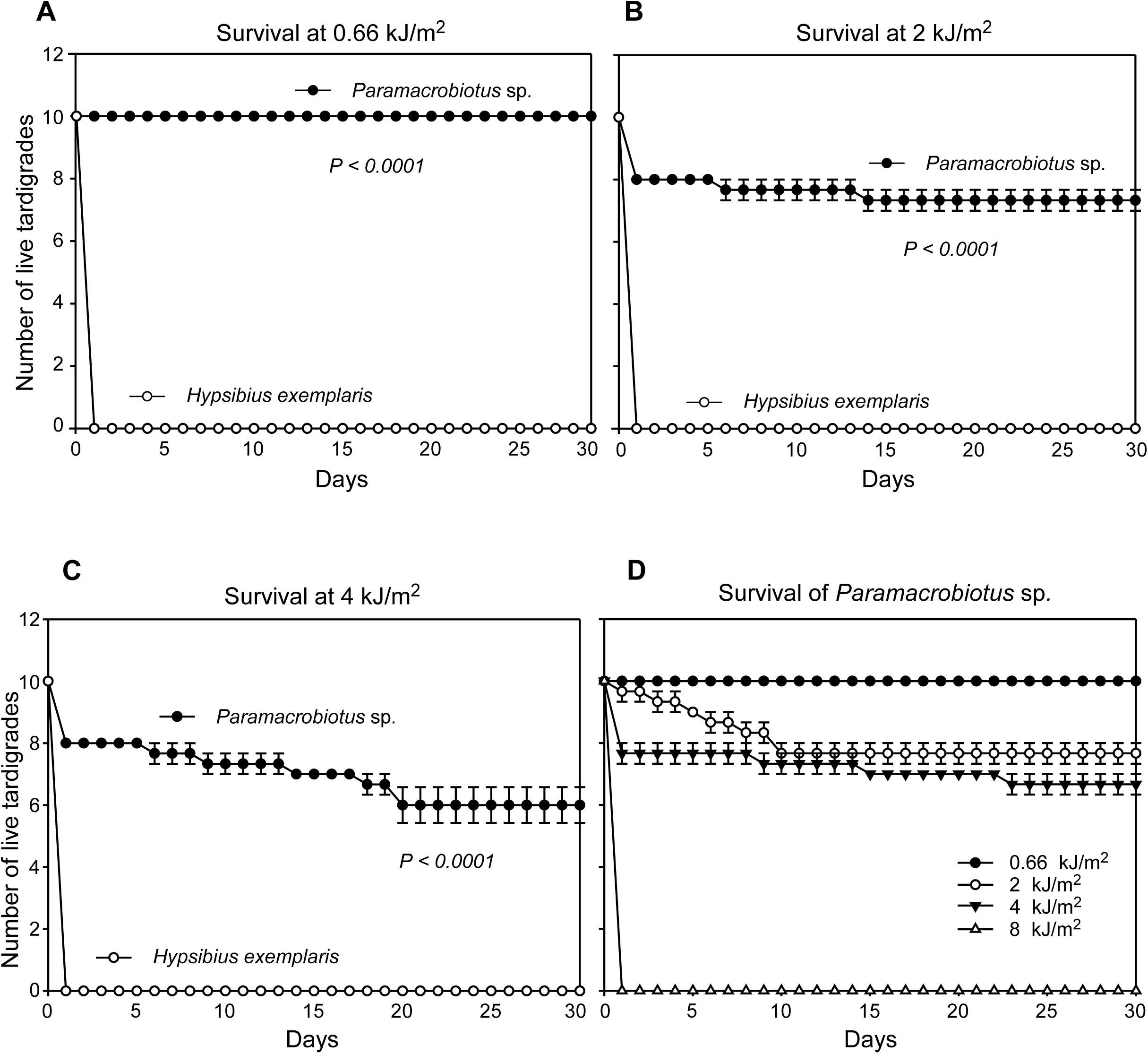
*Paramacrobiotus* sp. shows tolerance to UV radiation. Survival of *Paramacrobiotus* sp. under UV radiation exposure for 10 min (0.66 kJ/m^2^) (A), for 30 min (2 kJ/m^2^) (B), and for 1 hour (4 kJ/m^2^) (C) is shown. *H. exemplaris*, another eutardigarde species, was used as control. Comparison of survival of *Paramacrobiotus* sp. to different doses of UV radiation is shown in (D). Each point in the graph shows mean ± SE, n=10×3. Statistical significance was calculated using log-rank test.

### Naturally occurring fluorescence of *Paramacrobiotus* sp. is essential for its tolerance to UV radiation

Incidentally, *Paramacrobiotus* sp. showed strong fluorescence under UV illumination (DAPI channel, Axio Observer.Z1). This fluorescence was absent in *H. exemplaris*, which were sensitive to UV radiation. Similar fluorescence was observed in the eggs of *Paramacrobiotus* sp., but not on its moulted cuticle (Fig 3A). Extract from *Paramacrobiotus* sp. obtained after homogenizing the organisms in tissue lysis buffer also showed strong fluorescence under UV illumination (254 nm and 365 nm). The fluorescence was absent in the extract from *H. exemplaris*. The fluorescence was intact even after proteinase K treatment of the lysate for one hour suggesting that the fluorescent compound is not a protein (Fig 3B).

**Figure 3.**
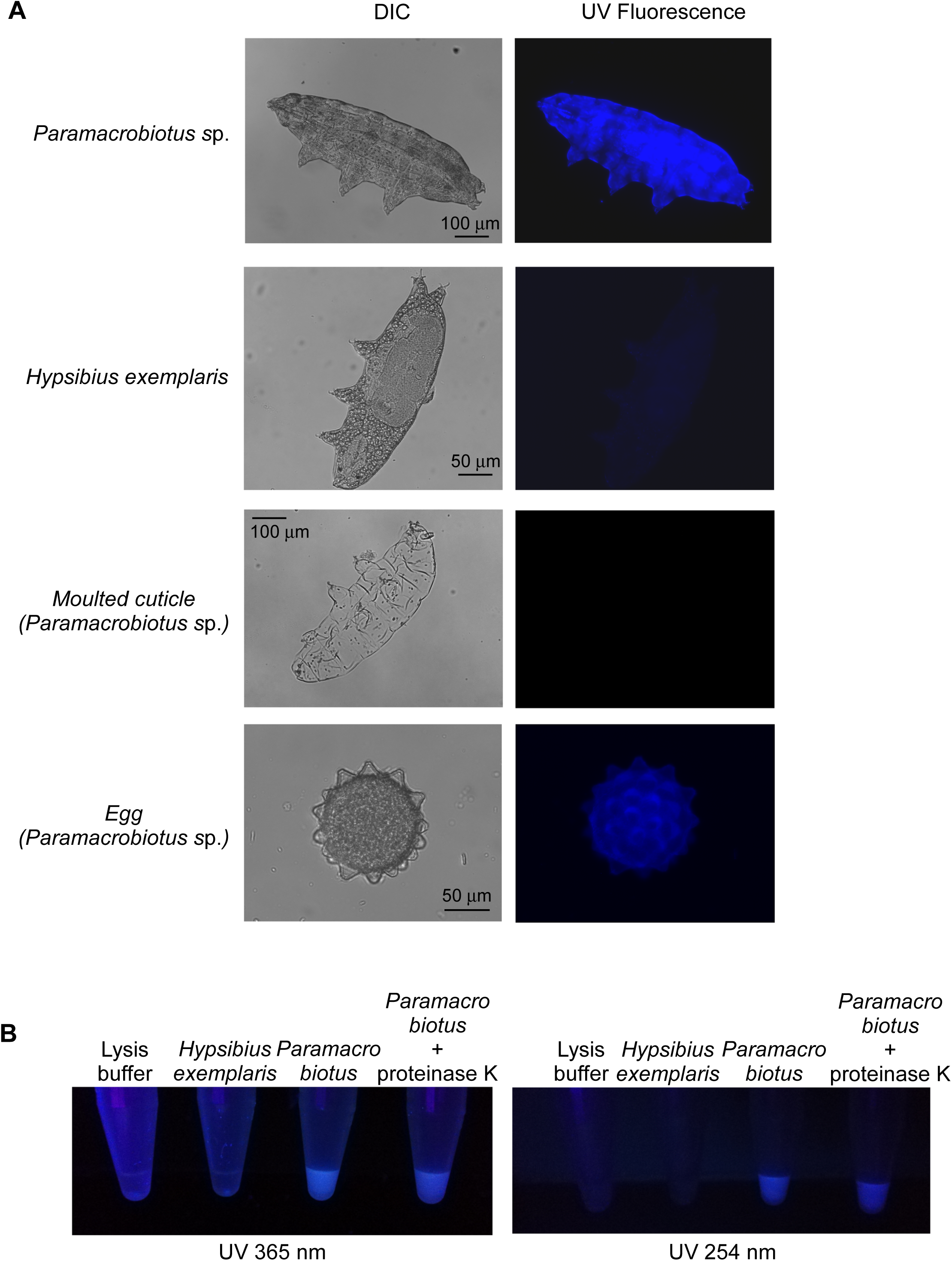
*Paramacrobiotus* sp. shows fluorescence under UV light. (A) Fluorescence microscope images of *Paramacrobiotus* sp., its moulted cuticle and egg are shown. UV-sensitive *H. exemplaris* was used as control for comparison. All images were taken under identical microscope settings. (B) Fluorescence images of the total lysates of *Paramacrobiotus* sp. and *H. exemplaris* under UV light (365 nm and 254 nm) are shown. *Paramacrobiotus* sp. lysate shows fluorescence that is stable even after proteinase K treatment.

Occasionally we come across hypopigmented *Paramacrobiotus* sp. during isolation. Morphological features and the nucleotide sequence of *ITS2* region revealed that the hypopigmented and the pigmented tardigrades belong to the same species (Figure 4A and S1). Interestingly, hypopigmented *Paramacrobiotus* sp. showed much less fluorescence under UV light (Figure 4A and B). When they were exposed to UV radiation for one hour, hypopigmented tardigrades showed significantly less UV tolerance compared to the pigmented ones. All hypopigmented *Paramacrobiotus* sp. died within 20 days after UV exposure, whereas 60% of the pigmented *Paramacrobiotus* sp. survived beyond 30 days (Fig 4C). This observation suggests that the newly identified *Paramacrobiotus* sp. uses fluorescence as a mechanism to resist harmful UV radiation.

**Figure 4.**
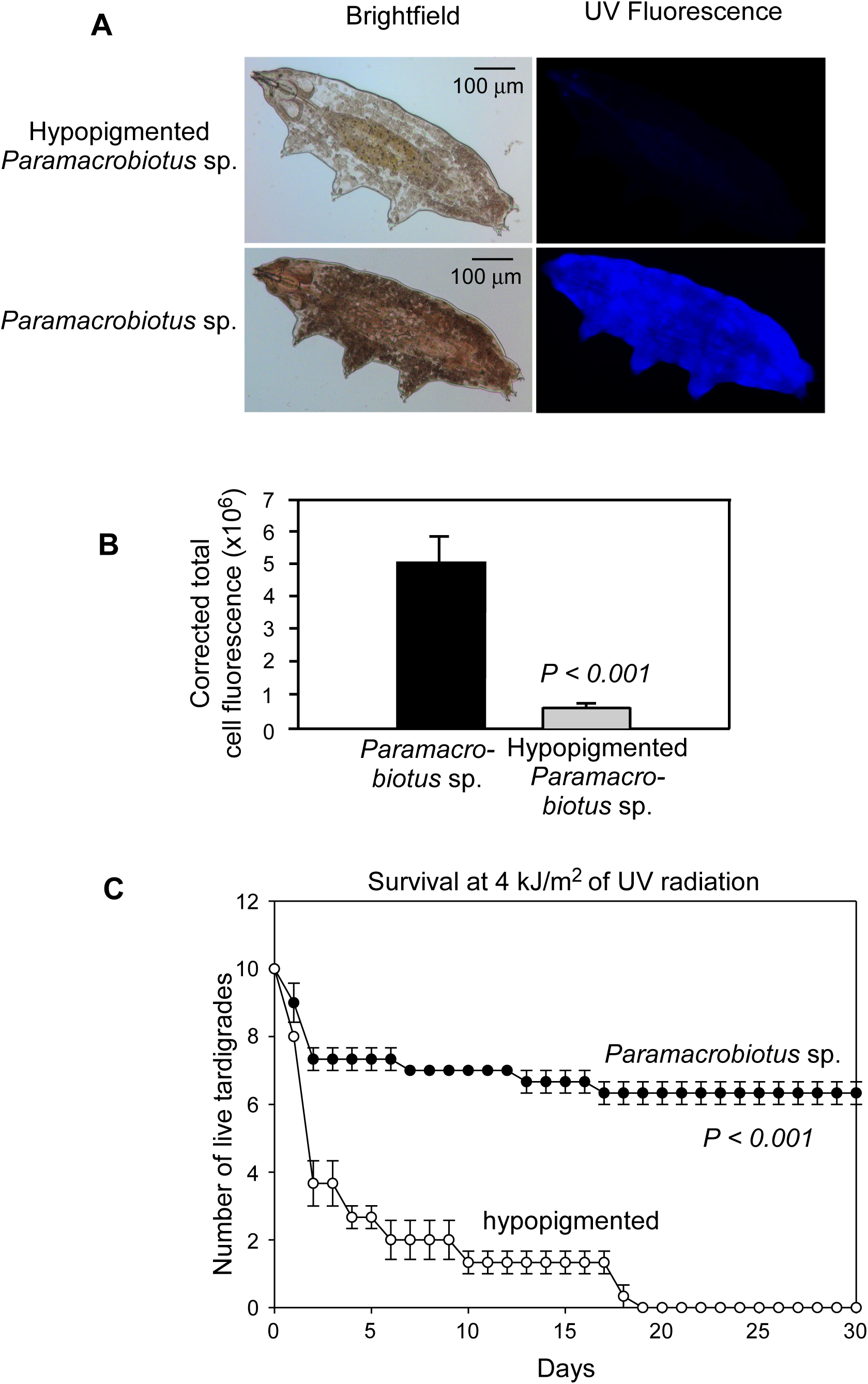
Hypopigmented *Paramacrobiotus* sp. does not show fluorescence and does not exhibit UV tolerance. (A) Images of hypopigmented *Paramacrobiotus* sp. showing reduced fluorescence under UV light compared to pigmented ones. (B) Graph showing the quantification of fluorescence. Bars show mean ± SE, n=6. Statistical significance was calculated using two-tailed Student’s t-test. (C) Survival of hypopigmented *Paramacrobiotus* sp. under UV radiation exposure for 1 hour (4 kJ/m^2^). Each point in the graph shows mean ± SE, n=10×3. Statistical significance was calculated using log-rank test.

### UV tolerance property of *Paramacrobiotus* sp. can be transferred to UV sensitive *H. exemplaris* and *C. elegans*

We tested if the UV tolerance can be transferred to UV sensitive *H. exemplaris.* For this, we homogenized 300 *Paramacrobiotus* sp. tardigrades in water. The supernatant of the homogenate was fluorescent under UV light (Fig 5A). UV sensitive *H. exemplaris* were covered in this fluorescent extract and exposed to UV radiation for 15 min (corresponds to 1 kJ/m^2^). *H. exemplaris* covered in water were used as control (Fig 5B). Interestingly, *H. exemplaris* tardigrades covered in the fluorescent extract showed partial tolerance to UV radiation. To further confirm the role of fluorescence, we photobleached the aqueous extract of *Paramacrobiotus* sp. by exposing it to 254 nm UV light. Bleached sample that lacked fluorescence was used for the experiment described above. Unlike the fluorescent extract of *Paramacrobiotus* sp., the photobleached non-fluorescent extract from the same did not confer UV tolerance on *H. exemplaris*. Similarly, extracts from hypopigmented *Paramacrobiotus* sp., which showed much reduced fluorescence, also could not confer UV tolerance on *H. exemplaris* (Fig 5C and D). Remarkably, the fluorescent extract of *Paramacrobiotus* sp. could confer partial UV resistance on *C. elegans*, a nematode (Fig 5E). Together, these results demonstrate that the fluorescence of *Paramacrobiotus* sp. is responsible for its UV tolerance.

**Figure 5.**
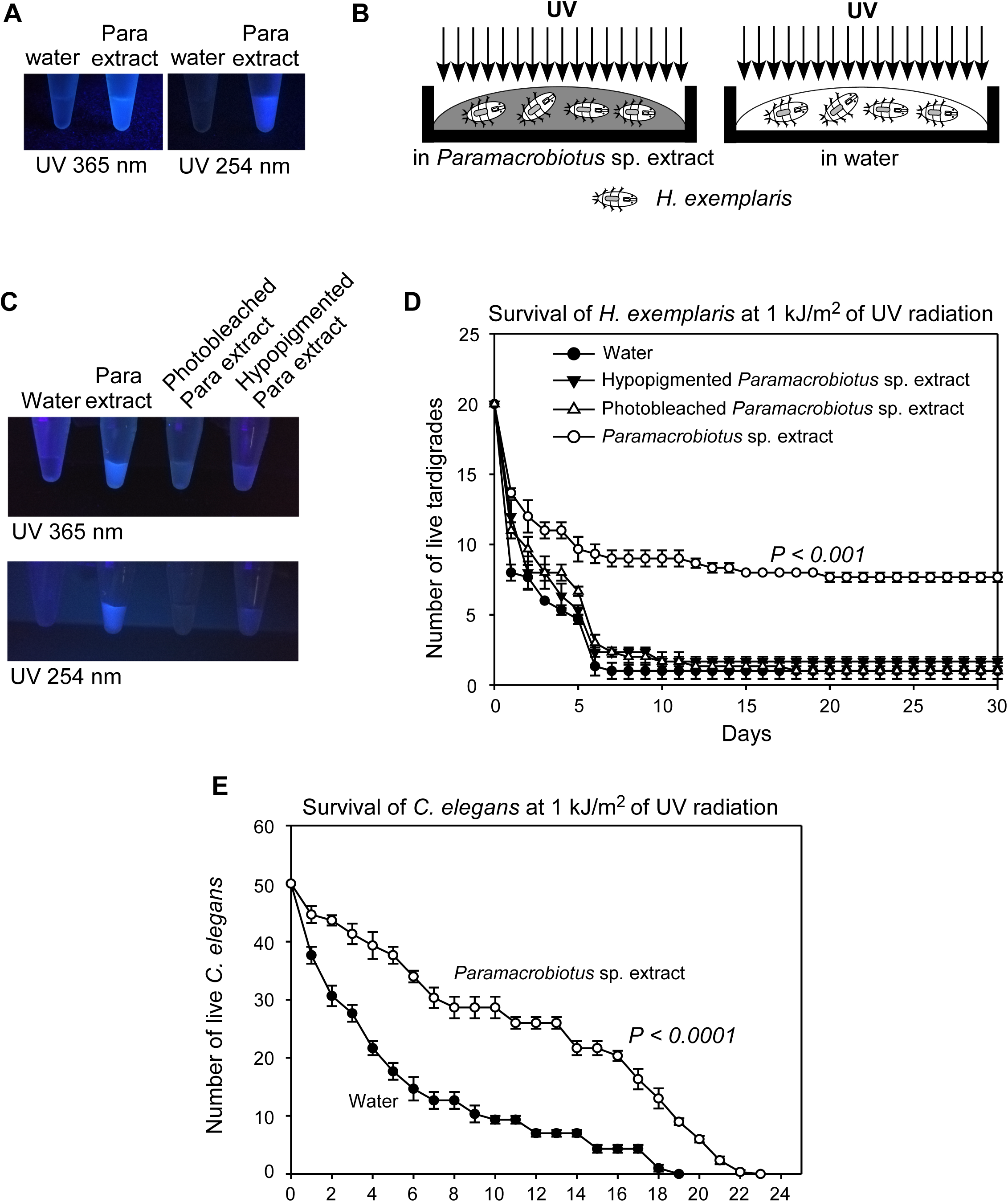
Transfer of UV tolerance property from *Paramacrobiotus* sp. to *H. exemplaris* and *C. elegans.*. (A) Fluorescence of aqueous extract from *Paramacrobiotus* sp. under UV light. (B) Schematic of the experimental setup. (C) Fluorescence of aqueous extract from *Paramacrobiotus* sp., photobleached extract and extract from hypopigmented strain. (D) Survival of *H. exemplaris* tardigrades incubated with extracts shown in (C) under UV radiation. (E) Survival of *C. elegans* incubated with fluorescent aqueous extract from *Paramacrobiotus* sp. under UV radiation. Each point in the graphs shown in (D) and (E) represents mean ± SE, n=20×3 (in D) or n=50×3 (in E). Statistical significance was calculated using log-rank test. Para, *Paramacrobiotus* sp.

### Properties of the fluorescent compound from *Paramacrobiotus* sp

We used methanol to extract the fluorescent compound from the newly identified tardigrade species. The methanolic extract from *Paramacrobiotus* sp. was fluorescent under UV light, whereas the extract from *H. exemplaris* was not (Fig 6A). We then subjected the methanolic extract of *Paramacrobiotus* sp. to High Performance Liquid Chromatography (HPLC) to isolate the fluorescent compound. We observed a unique peak (absorbance at 350 nm) in the extract from *Paramacrobiotus* sp. near 6 min, which was absent in the extract from UV-sensitive *H. exemplaris* (Fig 6B). The sample collected from this unique fraction of *Paramacrobiotus* sp. exhibited fluorescence under UV light confirming the isolation of fluorescent compound (Fig 6C). As expected, the fluorescent peak between 5 and 6 min in the HPLC profile of hypopigmented *Paramacrobiotus* sp. was much smaller compared to that in HPLC profile of pigmented *Paramacrobiotus* sp. (Fig 6D). Analysis using a spectrofluorometer showed that this fluorescent compound has excitation maxima (λ_ex_) at 370 nm and emission maxima (λ_em_) at 420-430 nm. Fluorescence was observed in a broad range of the UV spectrum between 250 to 370 nm (Fig 7A-C).

**Figure 6.**
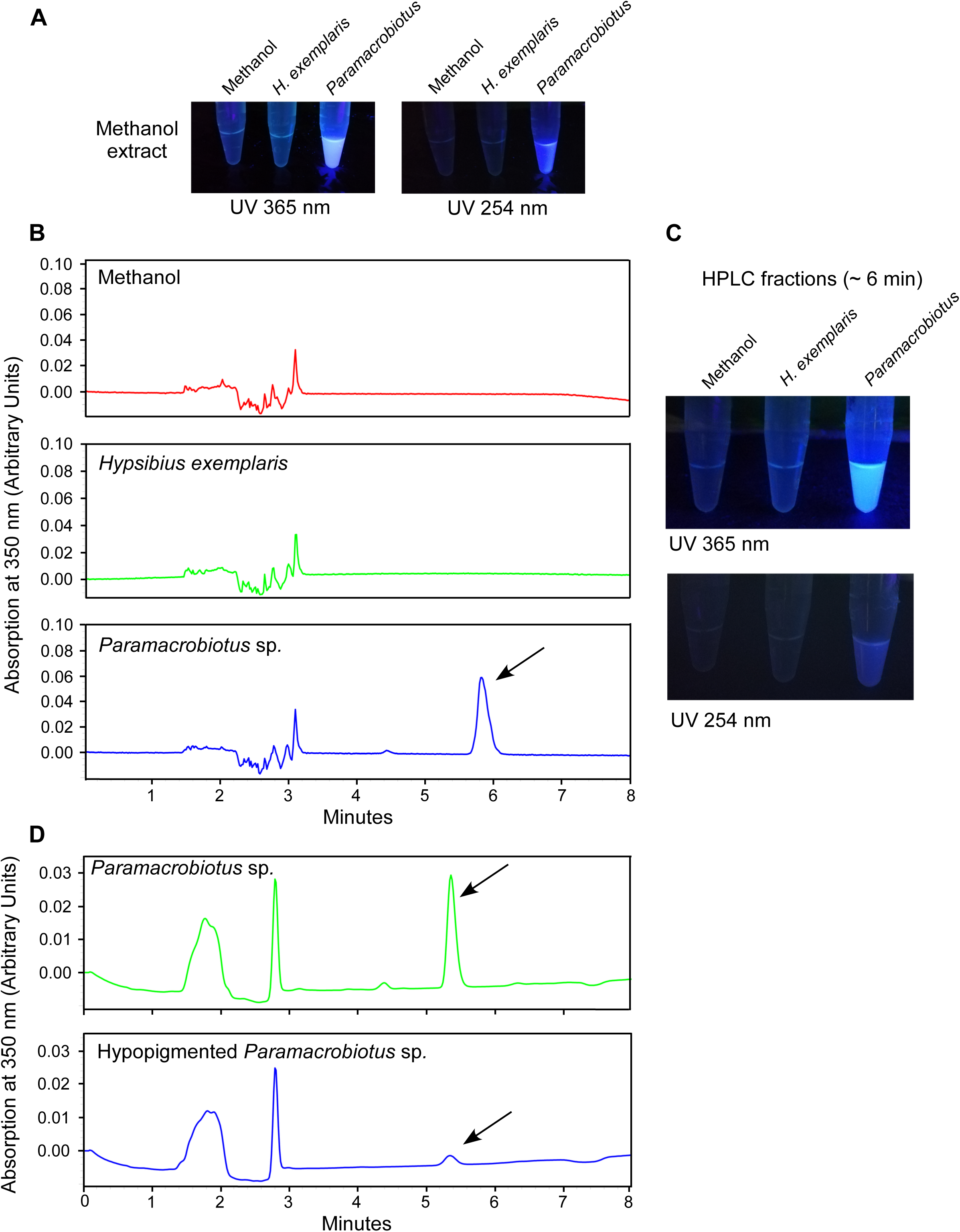
Isolation of fluorescent compound from *Paramacrobiotus* sp. by HPLC. (A) UV fluorescence of methanolic extracts from *Paramacrobiotus* sp. (B) HPLC profiles (absorbance at 350 nm) of methanolic extracts from *Paramacrobiotus* sp. and *H. exemplaris*. Arrow indicates the peak unique to *Paramacrobiotus* sp. UV fluorescence of the fraction from this peak is shown in (C). (D) HPLC profiles (absorbance at 350 nm) of methanolic extracts from hypopigmented *Paramacrobiotus* sp.

**Figure 7.**
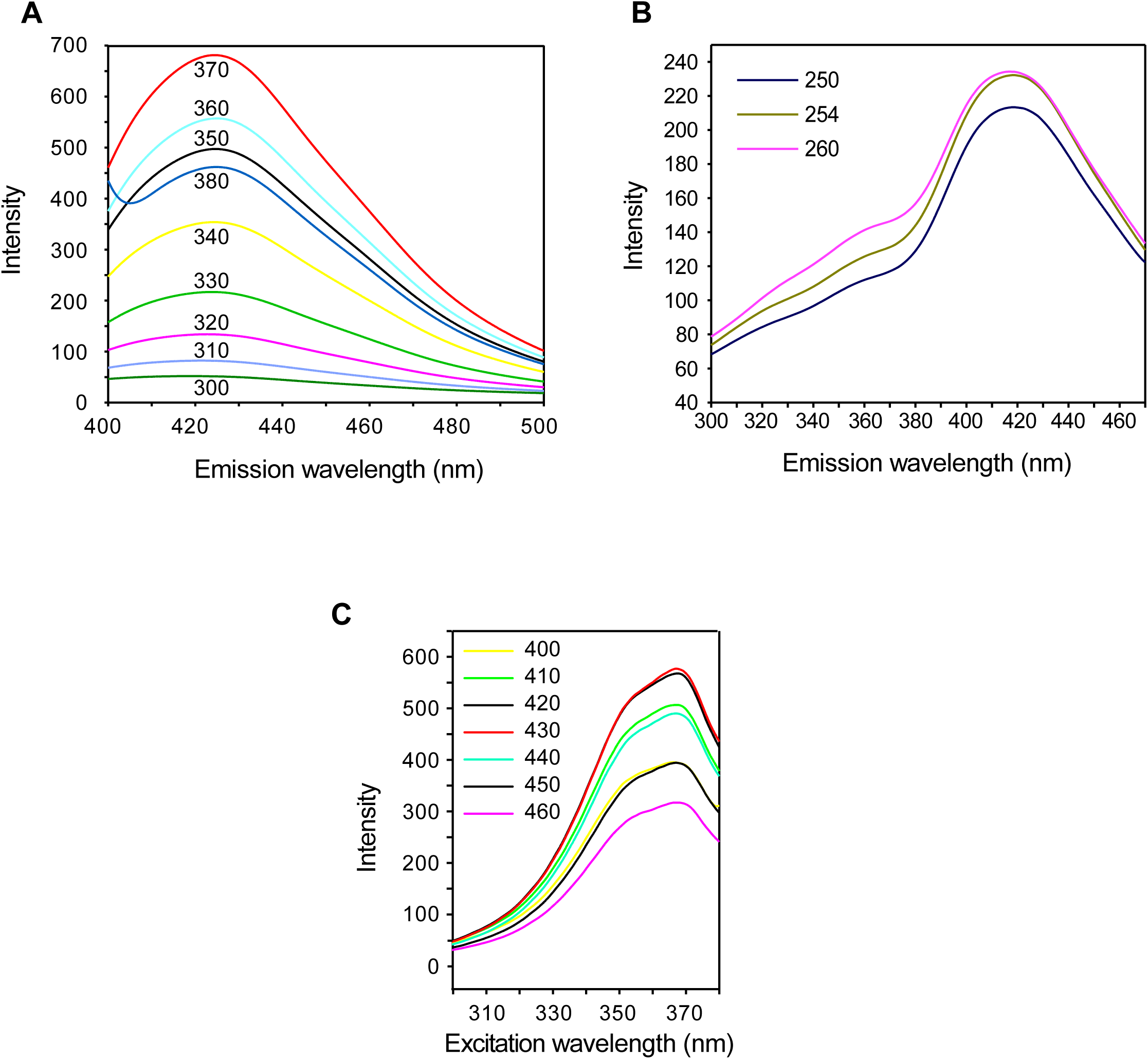
Spectral properties of fluorescent compound from *Paramacrobiotus* sp. Fluorescent HPLC fraction shown in Fig 6C was lyophilized and resuspended in methanol. Spectral property was analysed using spectrofluorometer. (A) Excitation scan: fluorescence intensity of the compound when excited using UV light of wavelengths ranging from 300 nm to 370 nm. (B) Relatively weaker fluorescence was seen at UV radiation of lower wavelength (250 nm, 254 nm and 260 nm). Two-times more concentrated extract was used for the analysis shown in (B) compared to the same in (A). (C) Emission scan: fluorescence intensity of the compound at multiple emission wavelengths (400 nm to 460 nm).

## Discussion

Fluorescence has been observed in diverse life forms - plants, worms, insects, frogs, birds, etc. (Lagorio et al., 2015). Functional significance of this phenomenon is known only in few organisms. Communication (e.g., mate attraction in zebra finch birds and luring preys in siphonophores) seems to be one of the primary physiological functions of fluorescence. However, the purpose of this phenomenon is not clear in most cases. Photoprotective role has been suggested for fluorescence in some organisms such as amphioxus, comb jellies and corals. In case of corals, a strong correlation has been demonstrated between fluorescence and the susceptibility to photo-inhibition and bleaching (Salih et al., 2000). However, there is no direct experimental demonstration of photoprotection imparted by fluorescence in any organism.

In this study, we show that a newly identified tardigrade species belonging to the genus *Paramacrobiotus* uses fluorescence to protect itself from UV radiation-induced death. A non-fluorescent variant of the same species was susceptible to UV radiation. Remarkably, we could transfer this property to a UV-sensitive tardigrade, *H. exemplaris*, and to *C. elegans*. Addition of fluorescent aqueous extract of *Paramacrobiotus* sp. on the surface of UV-sensitive *H. exemplaris* made it partially resistant to UV radiation. This was not observed when we used photobleached non-fluorescent extract or extract from hypopigmented *Paramacrobiotus* sp. These experiments demonstrate that fluorescence is responsible for UV tolerance in *Paramacrobiotus* sp. Thus, fluorescent pigment serves as a shield against UV radiation protecting these tardigrades from its lethal effects (Fig S2). *Paramacrobiotus* sp. has probably evolved this mechanism to counter high UV radiation of tropical southern India from where it was isolated. It is possible that these tardigrades have other mechanisms to protect themselves from UV radiation-induced damage. For example, robust DNA repair pathways. Analysis of its genome sequence will provide more insights.

Lethal effects of UV radiation are primarily due to DNA damage. It results in cyclobutane–pyrimidine dimers (CPDs) and 6–4 photoproducts (pyrimidine adducts) in genomic DNA, which affects replication and transcription. They can also cause lethal mutagenic effects (Sinha and Hader, 2002). UV radiation also damages DNA indirectly by producing reactive oxygen species. There are several organisms that resist lethal effects of UV radiation using multiple mechanisms. *Deinococcus radiodurans* has developed an efficient DNA repair pathway, which is responsible for its resistance to high ionizing radiation as well as UV radiation (Krisko and Radman, 2013). Production of pigments/compounds that absorb UV radiation is another mechanism commonly found in organisms from bacteria to mammals (Singh and Gabani, 2011). Cyanobacteria and other microbes produce UV-absorbing compounds such as scytonemin, myosporine and related amino acids (Oren and Gunde-Cimerman, 2007; Rastogi and Incharoensakdi, 2014). *Halobacterium salinarium*, a red pigmented bacterium, produces bacterioruberin which protects it from UV radiation (Dummer et al., 2011). Melanin in mammals and hipposudoric acid (red sweat) in hippopotamus are other examples of pigments that absorb UV radiation (Saikawa et al., 2004; Slominski et al., 2004). *Sander vitreus* (walleye) is a golden yellow fish found in North American lakes. Increased incidences of blue coloured walleye in recent years is thought to be an adaptation to increased UV radiation. Sandercyanin–Biliverdin complex is responsible for this blue colour. This complex exhibits red fluorescence under UV light (λ_ex_ 375 nm /λ_em_ 675 nm). However, it is not known whether this complex and its fluorescence protect the fishes from harmful effects of UV radiation (Ghosh et al., 2016). Our study adds fluorescence in the tardigrade *Paramacrobiotus* sp. to the list of UV-protection mechanisms. The chemical composition of this fluorescent compound remains to be investigated.

## Acknowledgements

We thank Prof. C. Jayabaskaran, Prof. Ganesh Nagaraju, Dr. Varsha Singh and Dr. Aravind Penmatsa for technical help.

